# Piezo1 stretch-activated channel activity differs between bone marrow-derived and cardiac tissue-resident macrophages

**DOI:** 10.1101/2023.11.27.568894

**Authors:** A Simon-Chica, A Klesen, R Emig, A Chan, D Grün, A Lother, I Hilgendorf, U Ravens, P Kohl, F Schneider-Warme, R Peyronnet

**Author notes:** Correspondence to Rémi Peyronnet and Franziska Schneider-Warme, Institute for Experimental Cardiovascular Medicine, Elsässer Straße 2Q, D-79110 Freiburg. These authors contributed equally to this work (A Simon-Chica, A Klesen).

## Abstract

Macrophages (MΦ) play pivotal roles in tissue homeostasis and repair. Their mechanical environment recently emerged as a key modulator of various cell functions, and MΦ mechanosensitivity is likely to be critical for cellular activity in particular in a rhythmically contracting organ such as the heart. MΦ, *in-vitro*-differentiated from bone marrow (MΦ_BM_), form a popular cell model for research. This study explores the activity of stretch-activated ion channels (SAC) in murine MΦ_BM_ and compares it to SAC activity in cardiac tissue-resident MΦ (MΦ_TR_). Our main findings are: i) MΦ_BM_ and MΦ_TR_ have stretch-induced currents, indicating expression of functional SAC at their plasma membrane; ii) the current profiles in MΦ_BM_ and in MΦ_TR_ show characteristics of cation non-selective SAC; iii) unlike in MΦ_BM_, Piezo1 ion channel activity at the plasma membrane of MΦ_TR_ is not detectable, neither by assessing electrophysiological activity using the patch clamp technique, nor by measuring cytosolic calcium concentration upon perfusion with Yoda1, a Piezo1 channel agonist. In mature scars after ventricular cryoablation, stretch-induced current characteristics of MΦ_TR_ are not significantly different compared to non-injured control tissue, even though scars are expected to contain a mix of pre-existing and circulation-recruited MΦ. This suggests that MΦ invading injured cardiac tissue either phenoconvert their mechanosensitivity from MΦ_BM_ to MΦ_TR_, or that the *in vitro* differentiation protocols used to obtain MΦ_BM_ generate cells that differ from MΦ recruited from the circulation during tissue repair *in vivo*. Further investigations will explore SAC identity in lineage-traced MΦ in scar tissue, and compare mechanosensitivity of circulating monocytes with that of MΦ_BM_.

**Key points:**

- MΦ_BM_ and MΦ_TR_ have stretch-induced currents, indicating expression of functional SAC at their plasma membrane;
- The current profiles in MΦ_BM_ and in MΦ_TR_ show characteristics of cation non-selective SAC;
- Unlike in MΦ_BM_, Piezo1 ion channel activity at the plasma membrane of MΦ_TR_ is not detectable

## 1. Introduction

Biophysical signals are omnipresent in biological processes. Among them, mechanical stimuli are known to regulate essential cell functions such as migration, differentiation and metabolism.(Emig, Zgierski-Johnston et al. 2021, Romani, Valcarcel-Jimenez et al. 2021) Virtually all cell types are mechanosensitive, *i.e*. they can sense and respond to mechanical signals. Accordingly, mechanosensitivity has been demonstrated for cells of the innate immune system, such as monocytes and macrophages (MΦ). MΦ form a heterogeneous cell population with important roles in tissue homeostasis and inflammatory responses. In many organs, including the heart, MΦ can be divided into (i) tissue-resident (MΦ_TR_) derived from embryonically established components of the myocardium, and (ii) MΦ recruited to the tissue from circulating blood (monocyte-derived MΦ: MΦ_MD_). In addition, experimental research has relied on MΦ, derived *in vitro* from bone marrow (MΦ_BM_) by culturing cells during exposure to pre-conditioned medium (more detail below).(Weischenfeldt & Porse, 2008) The various MΦ subtypes are characterized by distinct transcriptional profiles, protein expression patterns, and functions.(Heidt, Courties et al. 2014, Bajpai, Schneider et al. 2018, Chakarov, Lim et al. 2019)

Recent studies have highlighted that MΦ communicate with other cells not only *via* autocrine and paracrine signalling pathways, but that they can also sense, transmit and respond to electrical (Hulsmans 2017, Simon Chica 2022) and mechanical signals.(Lee, Du et al. 2022) Several MΦ functions are governed by ion channels,(Vicente, Escalada et al. 2003, Villalonga, David et al. 2010, Scheraga, Abraham et al. 2016) and MΦ mechanosensitivity has been shown to involve stretch-activated ion channels (SAC), a family of well-characterised mechanosensors. More specifically, the cation non-selective SAC Piezo1, which is ubiquitously expressed in cells across different species and involved in a plethora of biological processes,(Beech and Kalli 2019, Delmas, Parpaite et al. 2022) has been shown to regulate immune cell activity. In murine MΦ_BM_, genetic deletion of Piezo1 reduces inflammation and promotes wound healing.(Atcha, Jairaman et al. 2021) In addition, Piezo1-mediated calcium entry is essential for MΦ_BM_ stiffness sensing, modulating MΦ polarisation.(Atcha, Jairaman et al. 2021) Using implantable hydrogels mimicking the stiffness of fibrin clots, Forget *et al*. showed that stiffness-dependent recruitment of circulating monocytes into murine skeletal muscle *in vivo* requires Piezo1.(Forget, Gianni-Barrera et al. 2019) Piezo1 was also found to be involved in shear stress-induced monocyte activation, altering monocyte activity and contributing to valve inflammation.(Baratchi, Zaldivia et al. 2020) Recent evidence hints at a regulatory role of Piezo1 in the initiation and progression of chronic inflammatory diseases in myocardial fibrosis, atherosclerosis, pulmonary fibrosis, obesity, and diabetes.(Liu et al., 2022) Solis *et al*. further demonstrated that cyclic hydrostatic pressure changes can induce a pro-inflammatory gene expression profile in mouse monocytes *via* Piezo1, whereas mice lacking Piezo1 in myeloid cells show reduced pulmonary inflammation during bacterial infections or fibrotic autoinflammation.(Solis, Bielecki et al. 2019)

These findings highlight that Piezo1-expression and/or activity can modulate essential functions of both MΦ_BM_ and monocytes. However, there is limited knowledge about the mechanism underlying responses to mechanical stimulation of tissue-resident immune cells. This is of particular interest for MΦ_TR_ that are embedded in mechanically active tissue, such as the rhythmically contracting heart, in which MΦ are exposed to a wide range of mechanical stimuli, including stretch, compression, shear, torsion and bending. Insight into the mechanosensitivity of MΦ_TR_ is a prerequisite for a comprehensive understanding of their role in the maintenance of cardiac structure and function.(Hulsmans, Clauss et al. 2017, Nicolas-Avila, Lechuga-Vieco et al. 2020)

Here, we assess SAC expression and activity in murine cardiac MΦ_BM_ and MΦ_TR_. RNA sequencing reveals expression of RNA, coding for several mechanosensitive ion channels in MΦ_BM_ and MΦ_TR_. Using the patch-clamp technique, we observe stretch-induced currents in isolated MΦ_BM_ and MΦ_TR_, showing typical characteristics of cation non-selective SAC. Based on electrophysiological measurements, calcium imaging, and pharmacological interventions, we suggest that – unlike in MΦ_BM_ – the contribution of Piezo1 to stretch-activated currents in MΦ_TR_ is minor. This indicates that different mechanisms underlie mechanosensitivity of tissue-resident and *in-vitro*-differentiated MΦ.

## 2. Materials and methods

### 2.1 Experimental animals

All animal experiments were carried out according to the guidelines in the Directive 2010/63/EU of the European Parliament on the protection of animals used for scientific purposes; they were approved by the local authorities in Baden–Württemberg, Germany (Regierungspräsidium Freiburg, G20/01), and by the institution (X19/01R).

MΦ_TR_ were isolated from B6.129P2(Cg)-Cx_3_cr_1_^tm2.1(cre/ERT2)Litt^/WganJ mice (Cx_3_cr_1_^eYFP/+^; JAX stock #021160) wherein enhanced yellow fluorescent protein (eYFP) is expressed under the control of the fractalkine Cx_3_cr_1_ promoter, thus labelling mononuclear phagocytes, *i.e.* MΦ, monocytes, and dendritic cells.(Parkhurst et al., 2013) Experimental mice (both sexes) were 8–18 weeks old, and age-matched litter mates not expressing eYFP served as controls. MΦ_BM_ were obtained from WT mice (C57BL/6J) as described below.

### 2.2 Ventricular cryoablation

Left ventricular cryoinjury was applied to 12-week old Cx_3_cr_1_^eYFP/+^ mice. To this end, mice were anaesthetised by intraperitoneal injection of 80-100 µL of anesthesia solution (20 mg/mL Ketamine [Ketaset, Zoetis, Parsippany-Troy Hills, NJ, USA], 1.4 mg/mL Xylacin hydrochloride [0.12% Rompun, Bayer, Leverkusen, Germany], in 114 mM NaCl [0.67% m/V, B. Braun Melsungen, Melsungen, Germany]). After induction of deep anesthesia, eye ointment (Bepanthen containing 50 mg/mL dexpanthenol, Bayer, Leverkusen, Germany) was applied, 500 µL glucose solution (278 mM glucose = 5% (m/V), B. Braun Melsungen) was injected intraperitoneally, and 250 µL of analgesia solution (10 µg/mL buprenorphine [Temgesic, Indivior Inc., North Chesterfield, VA, USA] in 154 mM NaCl [0.9% m/V, B. Braun Melsungen]) was injected subcutaneously in the neck. Mice were shaved at the left side of the thorax (precordial region) and on the right leg. Mice were placed on a warming platform of a small animal physiology monitoring system (Harvard Apparatus, Holliston, MA, USA), and the front extremities were fixed with tape. A rectal thermometer was inserted to allow regulation of the heating platform to ensure a body temperature of 37°C. Thermometer and tail were fixed with tape. Mice were intubated for external ventilation (Kent Scientific, Torrington, CT, USA; 40% O_2_, 120 breathing cycles per minute). Isoflurane was applied at 5% until the animal stopped spontaneous respiratory movements, and then reduced to 2.0-2.5%. An infrared blood oximeter was attached to the right leg to follow hemoglobin oxygen saturation. The surgical field was disinfected using Softasept N (B. Braun Melsungen). Skin and muscles were cut along the 3^rd^ intercostal space, and a rib spreader was introduced. The pericardium was cut and the epicardial surface dry-blotted using a cellulose pad. A metal probe (stainless steel, 2.5-mm hexagon) pre-chilled in liquid nitrogen was applied for 8-10 s on the free left-ventricular mid-wall, avoiding major coronary vessels. After retraction of the probe, the time until the tissue regained deep-red colour was monitored (within 5 to 10 s). The rib spreader was removed and the thorax closed using a 6-0 silk suture around the 3^rd^ and 4^th^ rib (4-5 single knots). Before final closure, any remaining air was removed from the thorax using a small cannula. Isoflurane application was stopped and the skin was closed with a 6-0 silk suture. Once the mouse started breathing independently, intubation and fixation were terminated, and the mouse was transferred to a heated and oxygenated wake-up chamber. Analgesia was maintained for 72 h post-ablation by twice-daily subcutaneous injection of 250 µL of 10 µg/mL buprenorphine (in 154 mM NaCl; morning and late afternoon). During the night, buprenorphine was supplied *via* the drinking water (10 µg/mL buprenorphine [Subutex lingual tablets, Indivior] in 20 mM glucose solution). Terminal experiments were performed 3-4 weeks after ventricular cryoinjury.

### 2.3 MΦ isolation

MΦ_TR_ were isolated from murine hearts as previously described.(Fernandez, Kopton et al. 2021, Simon-Chica, Fernández et al. 2022) In short, mice were euthanised by cervical dislocation, hearts were swiftly excised and gently washed with 20 mL of cold phosphate buffered saline (PBS, Sigma Aldrich (in mM: KH_2_PO_4_ 136, NaCl 58, Na_2_HPO_4_-7H_2_O 268)). Thereafter, tissue was minced into small pieces and enzymatically digested for 50 min under gentle agitation with a mixture of 450 U/mL collagenase I, 125 U/mL collagenase XI, 60 U/mL DNase I, and 60 U/mL hyaluronidase (all Sigma-Aldrich, Munich, Germany) in 1 mL PBS at 37°C. Cells were re-suspended in fluorescence-activated cell sorting (FACS) buffer (PBS with 1% fetal calf serum + 0.1% bovine serum albumin; all Sigma Aldrich).

Cardiac MΦ_TR_ were FACS-purified based on cytosolic eYFP fluorescence using an Aria III cell sorter (BD Biosciences, Franklin Lakes, NJ, USA). The gating strategy was based on (a) forward scatter and side scatter signals to eliminate cell debris, (b) side scatter area and side scatter height for doublet discrimination and (c) eYFP intensity for purification of our cell population of interest. Cells from eYFP-negative littermates served as controls for determining background fluorescence. FlowJo software was used to analyze the FACS data.

### 2.4 Cell culture

FACS-purified MΦ_TR_ were seeded onto 15-mm glass coverslips in 24-well plates at a density of 20,000 cells/mL in 1 mL Dulbecco’s Modified Eagle Medium (DMEM, Thermo Fisher Scientific, Waltham, MA, USA) supplemented with 10% heat-inactivated fetal calf serum and 1% penicillin/streptomycin, (172 µM streptomycin and 200 µM penicillin, all Sigma-Aldrich). Non-adherent cells were removed by changing medium before functional experiments. Experiments were conducted 5-30 hours after cell isolation.

MΦ_BM_ were *in-vitro*-differentiated from bone marrow, harvested from murine femurs and tibiae. Epiphyses were removed and a 23G needle was inserted into one end. The bone marrow was flushed by injecting 2-3 mL of PBS per bone. Afterwards, cells were centrifuged (400×g for 5 min at 4°C) and cultured for 7 days in DMEM supplemented with 10% FBS, 20% L929-conditioned medium, 5% horse serum and 1% penicillin/streptomycin. L929-cell conditioned medium was obtained from the supernatant of L929 cells, which served as a source of recombinant MΦ colony stimulating factor. Preparation of L929-conditioned medium followed the protocol described in (Weischenfeldt & Porse, 2008). Briefly, 0.5·10^6^ L929 cells were plated in a 75-cm^2^ flask containing 55 mL of L929 medium (DMEM supplemented with 10% heat inactivated FCS and 1% penicillin/streptomycin) and maintained in an incubator containing 5% CO_2_ at 37°C. After 7 days, the supernatant was collected, filtered through a 0.22 µm syringe filter (Thermo Fisher Scientific), and used for preparation of MΦ_BM_ medium. Surplus supernatant not utilised immediately was stored at −20°C. After 7 days in culture, MΦ_BM_ were harvested using trypsin (Fisher Scientific) and seeded onto coverslips and maintained in an incubator containing 5% CO_2_ at 37°C for up to 30 hours and for further characterisation. Experiments were conducted within 5-30 hours after cell plating.

A human atrial fibroblast line that we characterised earlier,(Künzel, Rausch et al. 2020) was cultured in DMEM supplemented with 2 mM L-alanyl-L-glutamine (GlutaMAX, LifeTechnologies, Darmstadt, Germany), 10% fetal calf serum and 1% penicillin/streptomycin, and used as a positive control for pharmacological interventions targeting SAC.

### 2.5 Patch-clamp recordings and mechanical stimulation

The patch-clamp technique was used to assess the activity of SAC at the plasma membrane. Cell-attached patch-clamp recordings were performed using bath and pipette solutions previously described for characterizing Piezo1 channels.(Jakob, Klesen et al. 2021) The pipette solution contained (in mM): NaCl 150, KCl 5, CaCl_2_ 2, and HEPES 10; pH 7.4 with NaOH, 310 mOsm/L, while the bath solution contained (in mM): KCl 155, EGTA 5, MgCl_2_ 3, and HEPES 10; pH 7.2 with KOH, 310 mOsm/L (osmolality measured using a K-7400 osmometer, Knauer Wissenschaftliche Geräte, Berlin, Germany). Cells were washed twice with the bath solution prior to recording and at least 5 min were allowed between culture medium removal and the first recordings to avoid any potential blocking effects of antibiotics on SAC.(Gannier, White et al. 1994) Membrane patches were stimulated with brief (500 ms) negative pressure pulses, applied *via* the patch pipette (average pipette resistance 1.2 MΩ) with 10 mmHg increments from 0 to −80 mmHg, using a pressure-clamp device (ALA High Speed Pressure Clamp-1 system; ALA Scientific, Farmingdale, NY, USA). Experiments were performed at room temperature (20°C), using a patch-clamp amplifier (200B, Axon Instruments, San Jose, CA, USA) and a Digidata 1440A interface (Axon Instruments). Currents were digitised at 3 kHz, low-pass filtered at 1 kHz, and analysed with pCLAMP10.3 (Axon Instruments) and OriginPro 2019 software (OriginLabCorporation, Northampton, MA, USA).

### 2.6 Ca^2+^ imaging and signal analysis

Prior to imaging, cells were washed in Ca^2+^-free PBS and incubated for 20 min at room temperature with 3 μM Fluo-4-AM (F14201, Thermo Fisher Scientific) in PBS while being protected from ambient light. From this step onwards, cells were no longer exposed to antibiotics. After 20 min, the loading solution was removed, cells were washed with PBS, and used for Ca^2+^ imaging within 5 to 10 min after exposure to modified Tyrode solution (in mM): NaCl 140, KCl 5.4, Hepes 10, Glucose 10, CaCl_2_ 1.8, MgCl_2_ 1.0; pH 7.4 with NaOH, 300 mOsm/L.

Ca^2+^ dynamics in MΦ were imaged on an inverted confocal microscope (TCS SP8 X, Leica Microsystems, Wetzlar, Germany) using a 40x water immersion objective (HC PL APO CS2 40x/1.10). Fluo-4-AM was excited using a 488 nm laser line and the fluorescence signal was recorded with a hybrid detector operated in photon counting mode. The detection window was set from 497 nm to 568 nm. In order to maximise signal intensity, the pinhole was opened (7.33 Airy units). The Piezo1 channel activator Yoda1 (5, 10, 20 or 30 μM, dissolved in 10 mM DMSOt, Tocris Bioscience, Bristol, UK) and the Transient Receptor Potential Vanilloid (TRPV4) agonist GSK1016790A (300 nM [Sigma-Aldrich] dissolved in modified Tyrode solution) were applied using a local perfusion system. A vacuum-based suction system maintained a constant solution volume in the dish (∼1.5 mL). Flow rate was set to 1 mL min^−1^ by adjusting the height of reservoirs for gravity-driven flow. The control solution for Yoda1 experiments contained DMSO equivalent to largest Yoda1 concentration tested (both for patch-clamp and Ca^2+^ experiments).

Ca^2+^ signals were analysed with ImageJ. The StarDist plugin(Schmidt, Weigert et al. 2018) was used for cell area detection in images. Thereafter, mean pixel intensities were obtained for each time point to quantify temporal responses to Yoda1 and GSK1016790A. Data was exported to Matlab for further processing. Following background subtraction and signal normalization, baseline fluorescence was measured as the average of the last 10 frames before application of Yoda1 and GSK1016790A. To determine the maximal amplitude of responses to different treatments, the peak fluorescence signal reached during treatment exposure was selected for each cell.

### 2.7 RNA sequencing: expression of genes coding for mechanosensitive channels

The mRNA expression of mechanosensitive ion channels was assessed using published RNA sequencing data of murine MΦ_BM_ and MΦ_TR_.(Simon-Chica, Fernández et al. 2022, Xu, Gao et al. 2023) Raw data sets on MΦ_BM_ were obtained from the National Center for Biotechnology Information’s sequence read archive (Gene Expression Omnibus accession numbers: GSM6506821, GSM6506822 and GSM6506823). Raw data on MΦ_TR_ were obtained from our previously published work.(Simon-Chica, Fernández et al. 2022) All data were analysed using the Galaxy platform.(Afgan, Baker et al. 2018) This involved pre-processing and mapping onto the murine reference genome (NCBI37/mm9) as previously described.(Simon-Chica, Fernández et al. 2022) Gene expression in both datasets was assessed using featureCounts.(Liao, Smyth et al. 2014) The results were transformed into transcripts per kilobase million (TPM) taking into account gene length and sequencing depth. TPM of selected SAC were used as a measure of their expression at the mRNA level.

### 2.8 Statistical analysis

Data distribution was assessed with the Shapiro–Wilk normality test. Normally distributed data were compared using Student’s *t*-test, and not normally distributed data were compared using the Mann-Whitney test. Data are expressed as mean±SEM, individual data points are shown to illustrate the variability, and P-values <0.05 are considered as indicative of statistically significant differences between means (*P<0.05; **P<0.01, ***P<0.001). Throughout the study, “N” refers to the number of hearts and “n” to the number of cells.

## 3. Results

### 3.1 Stretch induces inward currents in MΦ_BM_ and MΦ_TR_

We compared the mechanical responses of murine freshly isolated ventricular cardiac MΦ_TR_ to MΦ_BM_. To assess the presence of stretch-induced currents, we performed cell-attached patch-clamp recordings, while steps of negative pressure were applied *via* the recording pipette to locally stretch the cell membrane (Fig. 1A). We found stretch-induced inward currents in 67% of murine MΦ_BM_ (N=4, n=40/60) and 56% of MΦ_TR_ (N=6, n=30/54; Fig. 1B). Pressure steps swiftly activated currents with no pronounced inactivation and slow deactivation. Quantification of the average currents (mean current calculated over the duration of the pulse of pressure) across the patched membrane as a function of applied pressure is shown in Fig. 1C. Average currents measured during maximum pressures pulse were 17.7 ± 4.9 pA and 10.3 ± 2.2 pA for mouse MΦ_BM_ and MΦ_TR_, respectively. Apart from pressure pulses at −30 and −80 mmHg, no significant differences between stretch-induced currents from MΦ_BM_ and MΦ_TR_ are observed. Taken together, the observed inward currents upon pressure application indicate the presence of functional SAC in plasma membranes of MΦ_BM_ and MΦ_TR_.

**Figure 1:**
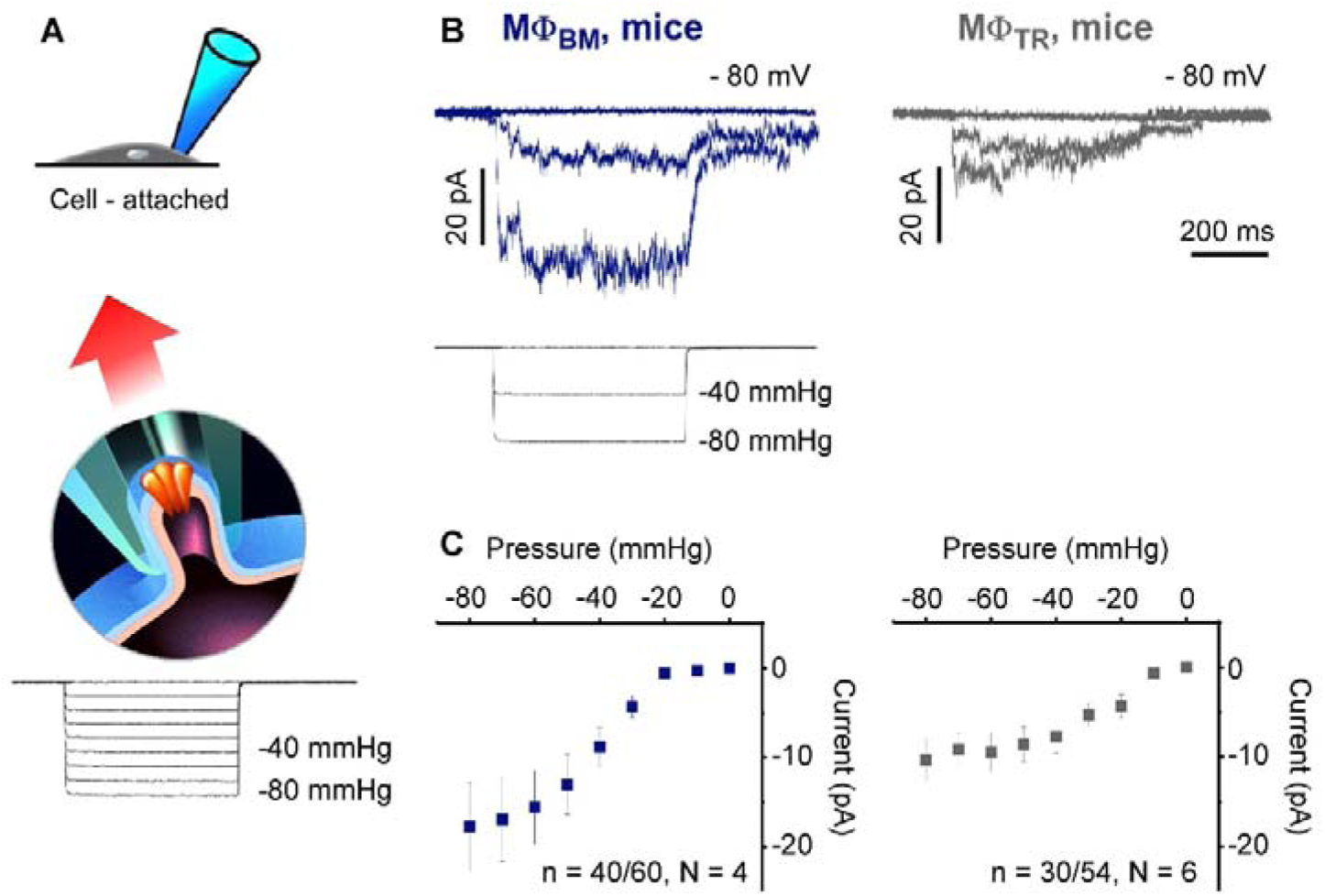
Stretch-activated channels (SAC) in MΦ from various species. (A) Cell-attached patch-clamp diagram used to investigate the presence of SAC (adapted from F. Aguila (CNRS), with permission). Bottom trace illustrates the pressure pulse protocol applied from 0 to −80 mmHg in 10-mmHg steps. (B) Representative traces of currents elicited at 0, −40 and −80 mmHg (as illustrated in bottom trace pulse protocol) in murine bone marrow derived macrophages (MΦ_BM_, blue, n = 40/60 responsive cells, 4 independent experiments/animals) and cardiac tissue-resident macrophages (MΦ_TR_) (grey, n = 30/54 responsive cells, 6 independent experiments). Holding potential of −80 mV. (C) Current-pressure relationships at a holding potential of −80 mV in MΦ_BM_ isolated from pig (n = 6 cells), mice (n = 40 cells) and in MΦ_TR_ (n = 30 cells). Results are shown as means ± SEM.

### 3.2 Stretch-induced currents show characteristics of cation non-selective SAC

In a subset of recorded MΦ membrane patches, we were able to discern individual channel opening events. Representative current traces at −20 mmHg and −120 mV are depicted in Fig. 2A. For further characterisation, we assessed single-channel current amplitudes at holding potentials from −120 mV to −40 mV. Current-voltage curves were fitted by linear regression, with the slopes corresponding to single channel conductances of 38.3 ± 0.2 pS and 44.3 ± 0.1 pS for MΦ_BM_ and MΦ_TR_, respectively (p=0.73, Mann-Whitney test). The reversal potentials, obtained by linear extrapolation, were 0.5 mV (−14.3 to 22.5 mV, 95% confidence interval) and −10.7 mV (−22.0 to 4.8 mV) for MΦ_BM_ and MΦ_TR_, respectively. We also assessed current responses to repeat application of −80 mmHg pressure pulses and found a decline in current amplitudes during repetitive stimulation (Fig. 2B). Normalised current amplitudes decay mono-exponentially with a half-maximal reduction after 6 to 8 iterations of negative pressure application (Fig. 2C; MΦ_BM_ N=2, n=6, MΦ_TR_ N=2 n=5. This data supports the hypothesis that SAC present and active in both types of MΦ are cation non-selective.

**Figure 2:**
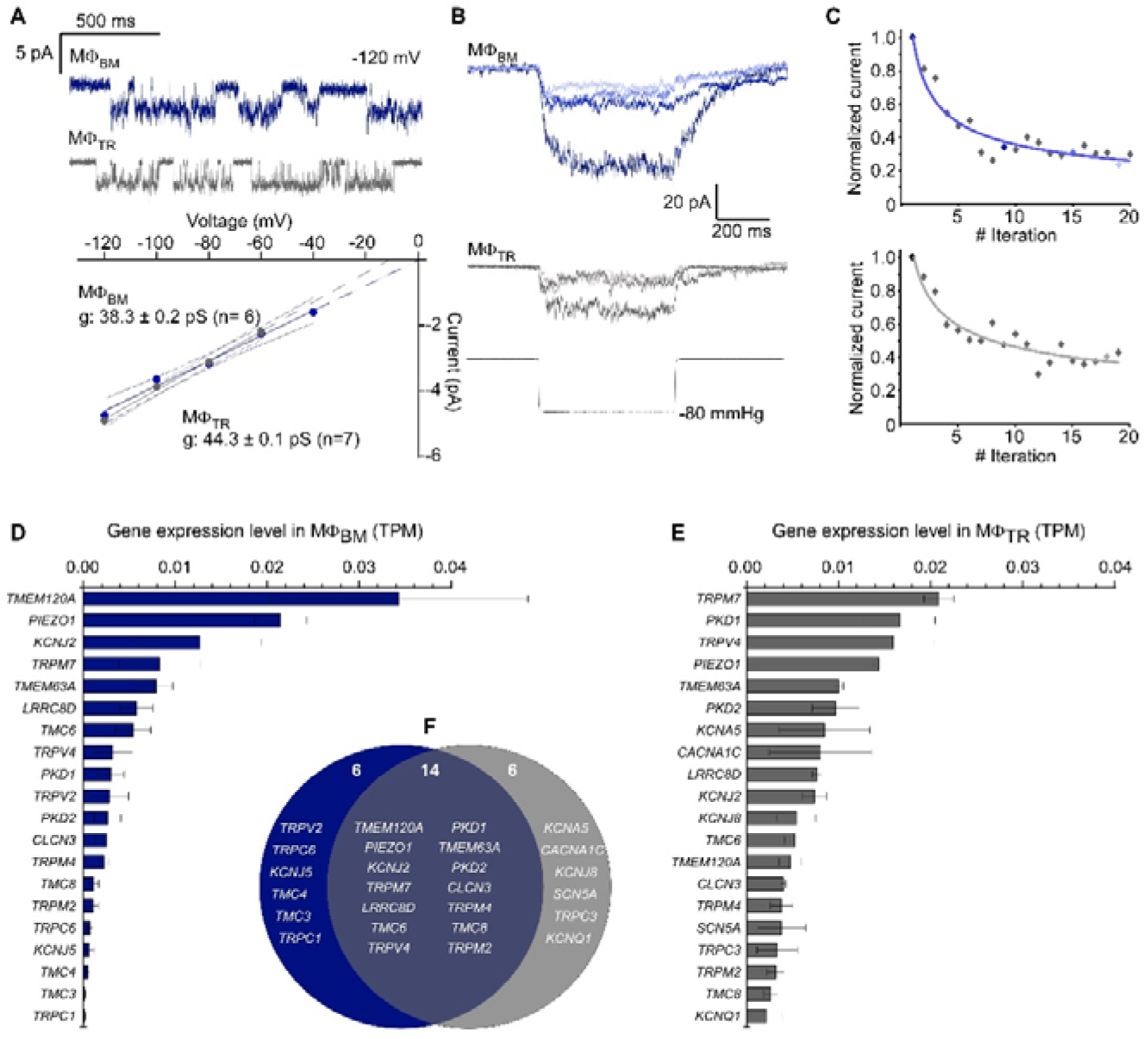
Biophysical characterisation of stretch activated channels (SAC). (A) Example traces of single channel activities from murine bone marrow derived macrophages (MΦ_BM_) in blue and tissue-resident macrophages (MΦ_TR_) in grey recorded at −20 mmHg at a holding potential of −120 mV (top). Current-voltage relationships for single channel currents. The slope of the linear regression gave the conductance. (B) Repetitive pressure stimulations (−80 mmHg) induced current ran-down at −80 mV in MΦ_BM_ (blue) and MΦ_TR_ (grey). (C) Normalised decreased mean current vs. number of pressure pulse iteration in MΦ_BM_ and MΦ_TR_. (D) RNA-seq data from MΦ_BM_ and MΦ_TR_ showing gene expression of genes coding for mechanosensitive channels.

### 3.3 MΦ express RNA coding for multiple SAC, including Piezo1

In order to identify molecular candidates that may underlie the observed stretch-induced currents, we analysed RNA expression levels in freshly isolated MΦ_TR_ and compared them to RNA expression data for MΦ_BM_. We focused on the expression of genes coding for mechanosensitive ion channels in the broadest sense, including ‘mechanically modulated’ (responding indirectly to mechanical stimulation or requiring co-activation) and ‘mechanically gated’ ion channels (changing their open probability in direct response to mechanical stimulation; see (Peyronnet, Nerbonne et al. 2016) for detail on nomenclature). We also included mechanosensitive channel candidates, *i.e*. ion conducting pathways that have not yet been functionally confirmed to be mechanosensitive, but which − based on phylogenetic analyses and sequence similarity, *e.g.* number of transmembrane domains of the corresponding proteins and/or links with mechanotransduction pathways – are predicted to be mechanosensitive. The 20 most highly expressed genes coding for mechanosensitive ion channels in MΦ_BM_ and MΦ_TR_ are presented Fig. 2D and E, respectively. Out of these, 14 genes were expressed in both MΦ_BM_ and MΦ_TR_ (Fig. 2F).

Based on the biophysical characterisation in section 3.2, the most plausible candidates underlying stretch-induced currents are genes coding for cation non-selective mechanosensitive ion channels. The most highly-expressed genes in MΦ_BM_ and MΦ_TR_ comprise *PIEZO1* (Fig. 2D, E and S1), several *TRP* channels from the melastatin (*TPRM7),* the polycystin (*TRPP1* and *2)*, and the vanilloid (*TRPV4)* families, as well as the recently identified SAC TACAN (Beaulieu-Laroche, Christin et al. 2020), *TMEM120A*; Fig. 2D and E).

### 3.4 Yoda1 increases stretch-induced currents in MΦ_BM_, but not in MΦ_TR_

As Piezo1 has been shown before to play a role in mechanosensation of murine monocytes and MΦ_BM_, we tested effects of the Piezo1 agonist Yoda1 on SAC activity. In MΦ_BM_, addition of 30 µM Yoda1 to the pipette solution resulted in a higher average stretch-induced currents (42.8 ± 6.3 pA, N=3, n=19; at −60 mmHg) compared to control conditions without Yoda1 (13.9 ± 2.2 pA, N=4, n=40; p<0.001, Mann-Whitney test, Fig. 3A). In contrast, stretch-induced currents recorded from MΦ_TR_ were not significantly affected by addition of 30 µM Yoda1 (9.9 ± 1.9 pA, N=4, n=24) compared to control recordings without the agonist (9.5 ± 1.8 pA, N=6, n=30; p=0.56, Mann-Whitney test, Fig. 3B).

**Figure 3:**
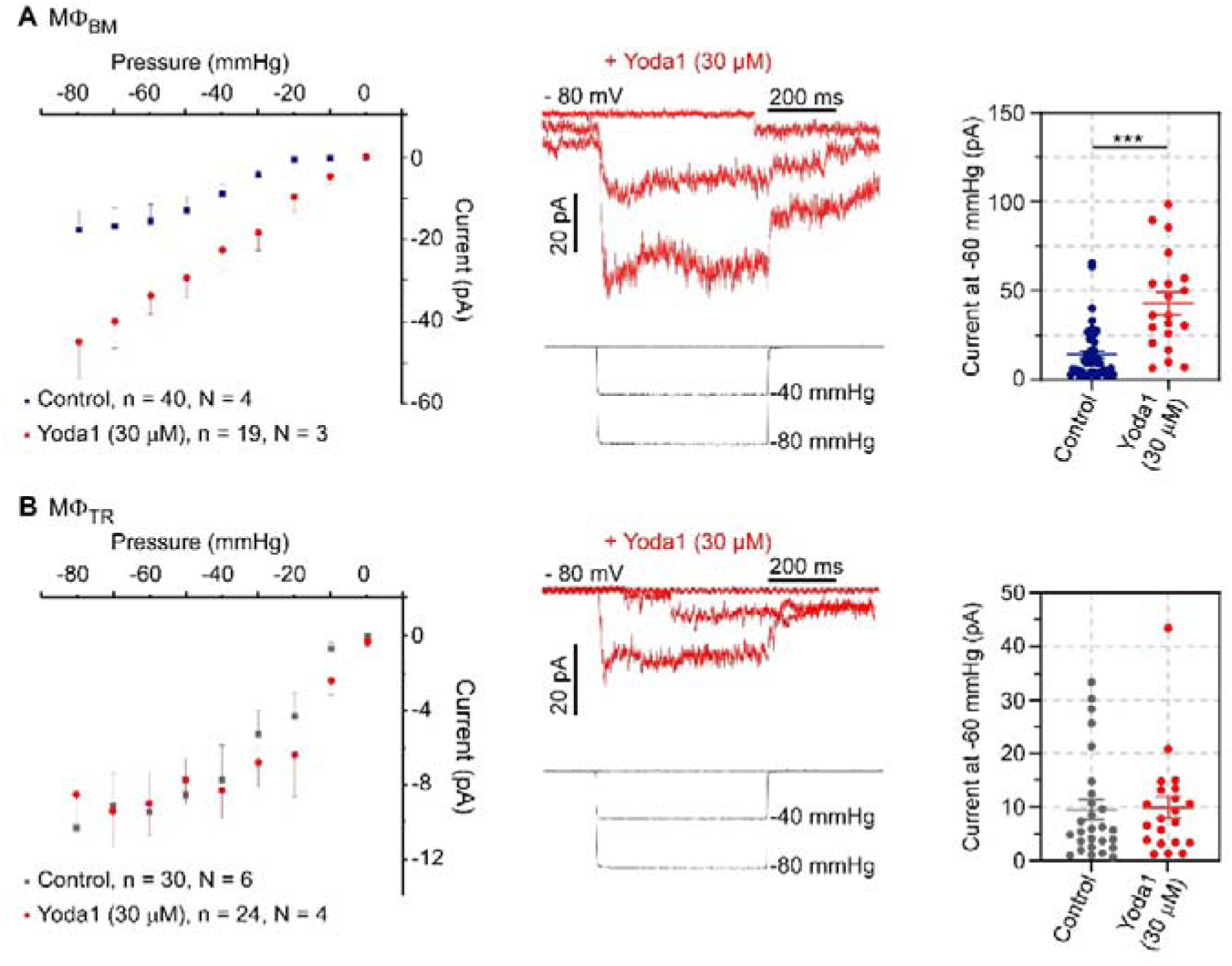
Yoda1 effect on stretch activated channel (SAC) activity. (A) Mean current amplitudes elicited by negative suction pulses in cell-attached patch-clamp mode in bone marrow derived macrophages (MΦ_BM_) in the absence (black, n = 40 cells) and presence of 30 μM (blue, n = 19 cells) Yoda1. (B) Mean current amplitudes elicited by negative suction pulses in tissue-resident macrophages (MΦ_TR_) in the absence (black, n = 30 cells) and presence of 30 μM (blue, n = 24 cells) Yoda1. Error bars indicate SEM. *P < 0.05, **P < 0.01, ***P < 0.001 by two-sample t test.

### 3.5 Yoda1 increases cytosolic Ca^2+^ concentration in MΦ_BM_ only

We monitored cytosolic Ca^2+^ concentration using the high-affinity fluorescent dye Fluo-4-AM as a complementary and more integrated readout to visualize potential Yoda-1 effects on Piezo1 activity at the whole-cell level and in a larger number of cells (Fig. 4A). Extracellular application of 5, 10 and 30 µM Yoda1 induced a significant increase in fluorescence of Fluo-4-loaded MΦ_BM_, indicating a rise in cytosolic Ca^2+^ concentration compared to control conditions (N=2, n=434, 446, 626 cells for 5, 10 and 30 μM Yoda1, respectively, and n=451 for DMSO-containing control; Fig. 4B). This Yoda1-induced increase in fluorescence was transient (Fig. 4C).

**Figure 4:**
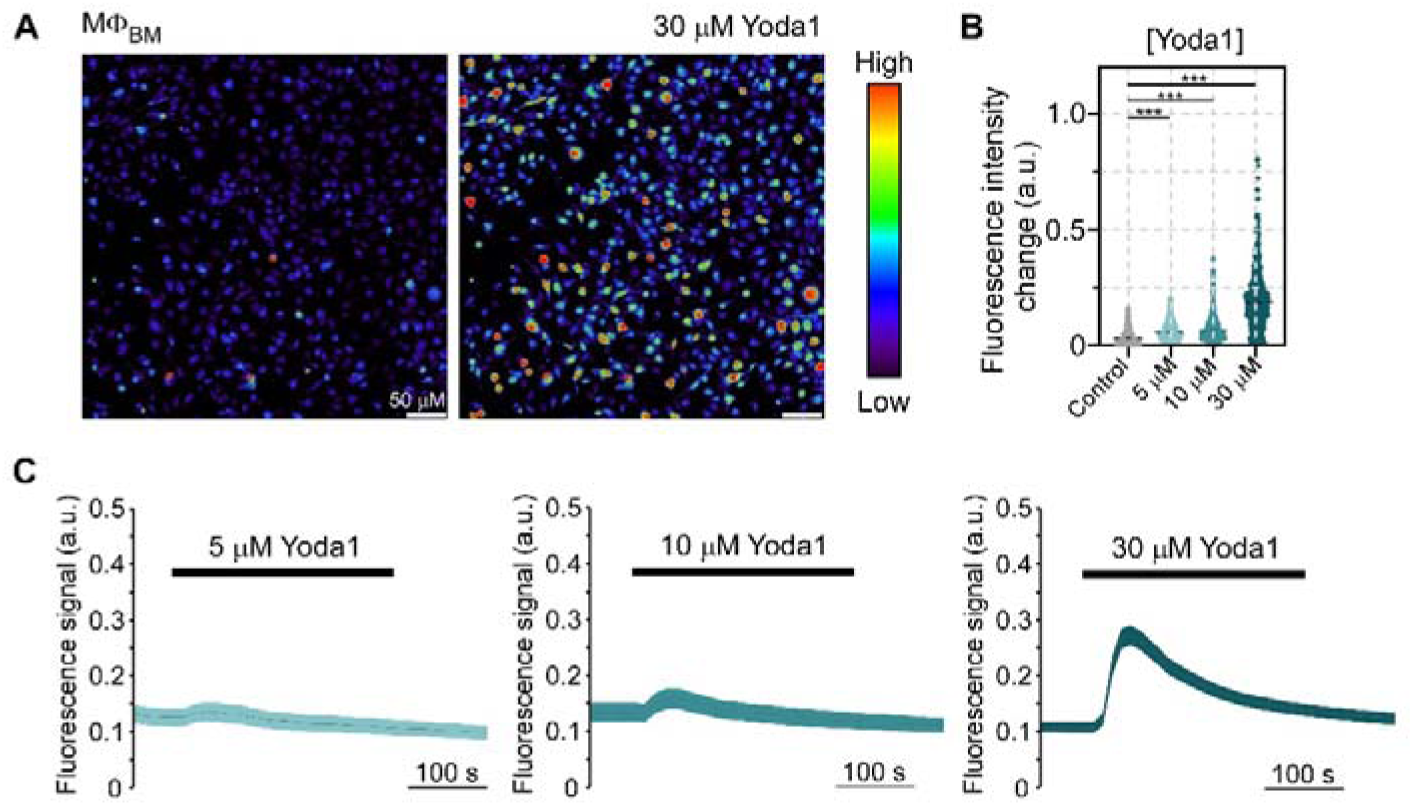
Cytosolic Ca^2+^ concentration in murine bone marrow derived macrophages (MΦ_BM_) in response to Yoda1 at different concentrations. (A) Representative images showing Ca^2+^ responses to 30 μM (peak intensity). (B) Peak intensity quantification per cell exposed to DMSO (N = 451 cells) or to increased concentration of Yoda1 (5, 10 and 30 μM. N = 434, 446, 626 cells, respectively). (C) Fluorescence signal traces averaged across all cells over time at 5, 10 and 30 μM Yoda1 concentration. Grey lines represent mean values and shaded regions SEM. *P < 0.05, **P < 0.01, ***P < 0.001 by two-sample t test.

In MΦ_TR_, we did not observe a significant increase in Fluo-4 fluorescence upon exposure to 20 µM Yoda1 (N=2, n=56; Fig. 5), while cardiac fibroblasts, known to express functional Piezo1 and used as an additional control, showed a robust response (Fig. S2). As the application of the Piezo1 agonist Yoda1 did not significantly affect stretch-induced currents or cytosolic Ca^2+^ concentration in MΦ_TR_, we tested an agonist targeting TRPV4 which, like Piezo1, is cation non-selective and strongly expressed in MΦ_TR_ (Fig. 2E). Application of the selective agonist GSK1016790A (300 µM) gave rise to a large and reversible increase in Fluo-4 fluorescence in MΦ_TR_ (N=2, n=77), indicating that TRPV4 is functionally active in MΦ_TR_.

**Figure 5:**
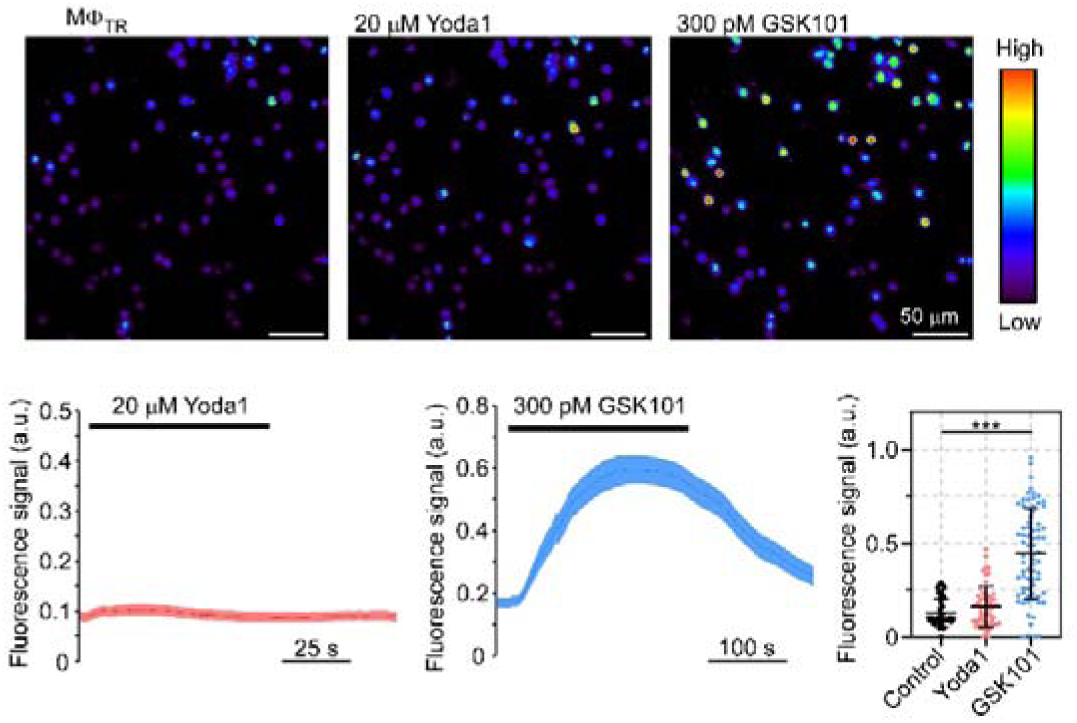
Regulation of Ca^2+^ influx in murine tissue resident macrophages (MΦ_TR_) upon application of Piezo1 and TRPV4 activators. Representative images showing Ca^2+^ responses to 20 μM Yoda1 and 300 μM GSK101 (peak intensity, top). Fluorescence signal traces averaged across all cells over time and quantification of peak intensities per cell (n=61, 56 and 73 cells exposed to control, 20 μM Yoda1 and 300 μM GSK101, respectively) (bottom). ***p < 0.001 (Student t-test).

### 3.6 Ventricular cryoablation does not change significantly stretch-induced current amplitude nor the number of Yoda1 responsive MΦ_TR_ compared to control

In the context of lesions, *e.g*. following myocardial infarction or ablation, monocyte-derived MΦ are recruited from the circulation into the myocardium. Accordingly we assessed Piezo1 activity in MΦ, isolated from murine hearts three to four weeks after ventricular cryoablation, which should contain a mix of MΦ_TR_ and MΦ_MD_. We assessed Yoda1 responses of stretch-induced currents in left ventricular MΦ from tissue that was remote to the scar (Fig. 6A) and in the scar (Fig. 6B), and compared this to left ventricular MΦ_TR_ isolated from sham operated animals (Fig. 6C). In all tested conditions, Yoda1 did not induce a significant increase in current amplitudes compared to control solution (Fig. 6D). In addition, no significant increase in the number of cells showing stretch-induced currents was observed upon Yoda1 application.

**Figure 6:**
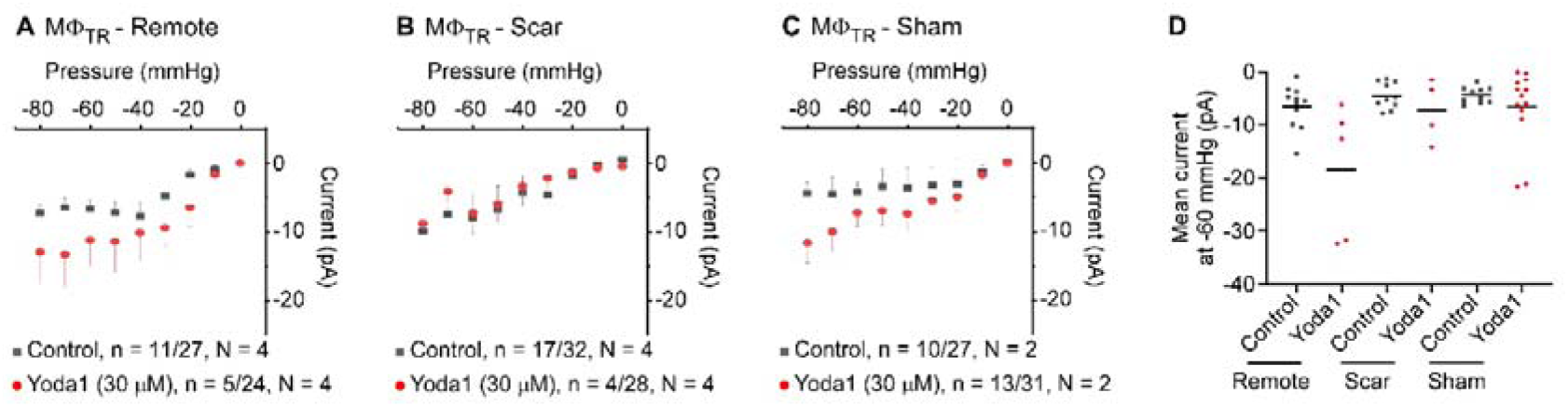
Yoda1 effect on and tissue-resident macrophages (MΦ_TR_) after cryoinjury. Current-pressure relationships at a holding potential of −80 mV in MΦ_TR_ isolated from regions remote to the scar (A), from the scar (B) and from sham operated animals (C)with or without Yoda1 (30 μM) in the pipette solution. (D) Current from individual cells at −60 mV. Under ‘n’, number of active cells (cells with SAC activity) over total number of cells recorded are shown respectively.

## 4. Discussion

MΦ contribute not only to maintenance of cardiac homeostasis, but also to tissue repair, such as during scar formation following myocardial injury. In the heart, MΦ_TR_ are exposed to rhythmically changing mechanical forces, based on the cardiac contraction-relaxation cycle. Additionally, the mechanical environment of MΦ_TR_ is altered during cardiac scar formation, associated with inhomogeneous tissue stiffening, which in turn changes regional tissue deformation, stretch and shear forces. In contrast to MΦ_BM_ and circulating monocytes, little is known about the ability of cardiac MΦ_TR_ to sense and respond to mechanical stimuli. This study compares the activity of SAC in mouse MΦ_BM_ to cardiac MΦ_TR_. Our main findings are: i) MΦ_BM_ and MΦ_TR_ have stretch-induced currents, indicating functional expression of SAC at their plasma membrane; ii) current profiles are comparable between MΦ_BM_ and MΦ_TR_, showing characteristics of cation non-selective SAC; iii) unlike in MΦ_BM_, Piezo1 ion channel activity at the plasma membrane of MΦ_TR_ is not detectable; and iv) stretch-induced current profiles are not significantly different in MΦ isolated from post-ablation tissue compared to MΦ isolated from remote or healthy control myocardium.

### 4.1 MΦ_BM_ and MΦ_TR_ functionally express SAC

We confirm the presence of stretch-induced currents in murine MΦ_BM_, and report similar current profiles in murine cardiac MΦ_TR_. Macroscopic current amplitudes (for the whole patch) do not differ significantly between the tested MΦ populations for pressure pulses from −30 mmHg to −80 mmHg, and current kinetics share fast activation, no or weak inactivation, and slow deactivation. Observed currents are inwardly directed at negative membrane voltages, with extrapolated reversal potentials in the region of −10 mV to 0 mV for MΦ_BM_ and MΦ_TR_. Together, these data confirm functional expression of cation non-selective SAC in the plasma membrane of MΦ_BM_ and MΦ_TR_. Additionally, we were able to record single channel activity in some membrane patches, and observed single channel conductances in the range of 38 to 44 pS for MΦ_BM_ and MΦ_TR_, respectively. The observed biophysical properties are shared by a number of mechanosensitive ion channels, including members of the TRP and Piezo families. This is in keeping with own and previously published RNA sequencing data, which shows that MΦ_BM_ and MΦ_TR_ express mRNA coding for TRPV, TRPM, TRPP and Piezo1 channels, albeit at different levels.

Direct comparisons of RNA expression data from MΦ_BM_ and MΦ_TR_ should be considered with caution, due to differences between protocols used to obtain the cells. We tried to control these differences in functional studies, and recorded all MΦ within the same time window after plating them (5-30 hours). That said, MΦ_BM_ were cultured and differentiated for 7 days before plating, while MΦ_TR_ were obtained acutely by enzymatic digestion, and then sorted (FACS) and plated.

### 4.2 Despite presence of mRNA transcripts, Piezo1 channel activity was not detected in MΦ_TR_

RNA sequencing data reveal that Piezo1 mRNA is expressed in MΦ_BM_ and MΦ_TR_, in keeping with another recent study.(Lother, Bondareva et al. 2021) However, while the application of the Piezo1-specific agonist Yoda1 increased stretch-induced currents, leading to a transient rise in the intracellular Ca^2+^ concentration in MΦ_BM_, Yoda1 application did affect neither current amplitudes nor Ca^2+^ concentrations in isolated cardiac MΦ_TR_. The apparent discrepancy between mRNA levels and functional experiments may be explained by Piezo1 localisation on intracellular membranes, rather than the plasma membrane. Such endomembrane localisation of Piezo1 has been described for the endoplasmic reticulum of epithelial cells,(McHugh, Buttery et al. 2010) and for the endoplasmic /sarcoplasmic reticulum, mitochondria, and nucleus of vascular smooth muscle cells.(Liao, Lu et al. 2021) As MΦ_TR_ form a heterogeneous population, Piezo1 could be expressed in a sub-population of MΦ_TR_ only that may not be dominant in the cultures used for this study. This would be in line with the observation in monocytes that Piezo1 is present only in a unique CD11b^+^/CD115^+^ subpopulation, representing about 25 – 40 % of the overall monocyte population under the conditions of the respective study.(Forget, Gianni-Barrera et al. 2019) However, our previous study (Lother, Bondareva et al. 2021) showed that Piezo1 is expressed both in CD45.2^+^ CD11b^+^ positive cells with high F4/80 and low Ly6C expression (macrophages), and in cells with low F4/80 and high Ly6C expression (monocytes), making this hypothesis unlikely. An alternative explanation of the observed discrepancy is that plasma membrane-localised channel proteins may be affected by the cell isolation procedure, including treatment with collagenase I and XI, which is necessary to obtain MΦ_TR_ but not MΦ_BM_. That said, the same cell isolation protocol was used in a previous study where levels of K^+^ channel-encoding mRNA matched structural observations and K^+^ channel activity in MΦ_TR_.(Simon-Chica, Fernández et al. 2022) In the case of Piezo1, lack of a reliable antibody (in our hands) for immunocytochemical analyses impedes validation of the putative subcellular location of the protein.

### 4.3 Yoda1 effects are different in patch-clamp recordings compared to Ca^2+^ imaging experiments

Currently, Yoda1 is the most commonly used agonist to pharmacologically activate both murine and human Piezo1 channels, affecting the sensitivity and inactivation kinetics of mechanically-induced currents.(Syeda, Xu et al. 2015) In the present study, we observe differential effects of Yoda1 in patch-clamp recordings compared to Ca^2+^ imaging experiments. In cell-attached patch-clamp recordings, addition of Yoda1 to the pipette solution does not induce large Piezo1 activity at baseline, but increases current amplitudes in the presence of additional mechanical stimulation (observations by us and others, and not limited to MΦ). Thus, in patch-clamp recordings, Yoda1 mainly sensitizes Piezo1 to stretch, rather than activating channel gating in control conditions. In contrast, bath perfusion with Yoda1 is sufficient to induce an increase in cytosolic Ca^2+^ concentration, as observed by fluorescence imaging. These differential effects of Yoda1 application are probably based on the different mechanical environments that the cell membrane exposed to Yoda1 faces in the different experiments. We performed Ca^2+^ imaging experiments to indirectly assess Ca^2+^ currents across the plasma membrane without local application of mechanical deformation. We used cell-attached recordings to track ion channel activity before, during and after defined levels of membrane stretch. In the cell-attached mode, recorded channels sit in a membrane patch surrounded by the small opening of the glass pipette, thereby creating a particular mechanical environment even before additional stretch application (inducing high membrane tension, changing membrane curvature, possibly shifting lipid and protein composition, and minimizing flow-based shear).(Suchyna, Markin et al. 2009) This particular mechanical state may change the reference level of Piezo1 activity at rest. Additionally, as highlighted above and elsewhere,(Liu, Hu et al. 2022) Ca^2+^ imaging may reveal also Piezo1-mediated Ca^2+^ release from intracellular stores, which is not detectable with patch-clamp recordings of plasma membrane activity. While the mode of action is different between the two types of experiments (patch-clamp *vs* Ca^2+^ imaging), Yoda1 consistently increased SAC activity in MΦ_BM_, but not in MΦ_TR_.

### 4.4 Piezo1 may be functionally coupled to secondary ion channels

RNA sequencing data shows expression of a wide array of mechanosensitive channels, including various TRP channels in MΦ_BM_ and MΦ_TR_. This is in line with previous studies reporting the expression and activity of several members of the TRP family (including TRPM2/4/7/8, TRPC1/3/6 [canonical], TRPV1/2/4, TRPA1 [ankyrin], TRPM1/2/3 [mucolipins], TRPP2) in myeloid cells. These channels have been implicated in the control of cell proliferation and survival, polarisation, mobility, phagocytosis, intercellular communication, and inflammatory responses (as reviewed in (Santoni, Morelli et al. 2018, Selezneva, Gibb et al. 2022)).

We find that pharmacological activation of TRPV4 leads to a strong increase in cytosolic Ca^2+^ in MΦ_TR_, in line with the recently reported involvement of TRPV4 in stretch sensing in cardiac MΦ_TR_,(Wong, Mohan et al. 2021) and in matrix stiffness sensing in skin MΦ.(Dutta, Goswami et al. 2020) TRP channels are not necessarily directly activated by stretch,(Nikolaev, Cox et al. 2019) but may respond to stretch-induced activation of Piezo1, leading to cytosolic Ca^2+^ entry and increased open probability of Ca^2+^-activated TRP channels.(Yu, Gong et al. 2022) Such functional coupling, involving the several TRP channel subtypes, may underlie the large Ca^2+^ entry upon Yoda1 application in MΦ_BM_. Secondary channel activation *via* Piezo-induced Ca^2+^ entry has also been shown for Ca^2+^-activated K^+^-selective channels, such as the large-conductance BK channels,(Jakob, Klesen et al. 2021) which is lowly expressed in the here studied MΦ populations (not shown). Further experiments are needed to dissect the functional interactions of SAC with one-another, both at the plasma membrane and at intracellular membranes, to fully understand channel-based MΦ mechanosensation.

### 4.5 MΦ_BM_ may not represent a suitable model to study SAC activity in MΦ_TR_ and MΦ_MD_

While MΦ_BM_ provide a convenient model to study MΦ properties, the extent to which these cells reflect MΦ_TR_ and MΦ_MD_ mechanosensitivity is unknown. We find that, in contrast to MΦ_BM_, Piezo1 activity is minor at the plasma membrane of MΦ_TR_ isolated from healthy myocardium. This suggests that MΦ_BM_ may not be a suitable model system for the study of MΦ_TR_ mechanosensitivity.

As Piezo1 expression is upregulated in pathologically remodelled tissue exhibiting increased matrix stiffness,(Chen, Wanggou et al. 2018, Jakob, Klesen et al. 2021, Li, Zhang et al. 2022) we explored whether Piezo1 activity in cardiac MΦ is altered in lesioned ventricular tissue. To this end, we compared stretch-induced currents and responses to Yoda1 application in murine MΦ, isolated from scar tissue and remote myocardium 4-weeks after ventricular cryoablation to those in control MΦ_TR_. Ventricular cryoinjury did not increase the percentage of cells responding to membrane stretch and did not result in significantly larger currents in the presence of Yoda1 even though the myocardium after ventricular injury is expected to contain both MΦ_TR_ and MΦ_MD_. This may suggest that MΦ_BM_ are not a suitable model system for the study of MΦ_MD_ mechanosensitivity either. Alternatively, Piezo1 activity, expression or/ and localisation in MΦ_MD_ may change over time in cardiac tissue, in line with reported transcriptional and phenotypic changes of MΦ during mature scar formation. This may shift MΦ_MD_ phenotype towards MΦ_TR_ characteristics at late time points after cardiac injury, as suggested before.(Mouton, DeLeon-Pennell et al. 2018, Walter, Alonso-Herranz et al. 2018, Dick, Macklin et al. 2019)

Taken together, these results suggest that the *in vitro* derived MΦ_BM_ model is not suitable for characterization of SAC activity in freshly isolated myocardial MΦ, potentially independent of their lineage. In future studies, it would be interesting to selectively characterise the SAC activity profile of MΦ populations present at defined time points following myocardial injury.

In conclusion, this study confirms the presence of cation non-selective SAC in MΦ_BM_ and characterises them in MΦ_TR_. While Piezo1 mediates stretch-induced currents in MΦ_BM_, Piezo1 activity at the plasma membrane is minor in MΦ_TR_, where TRPV4 channels appear to underlie SAC activity. Although MΦ populations in lesioned myocardium are expected to include cells derived from circulating monocytes, for which Piezo1 activity has been shown to be of functional importance,(Forget, Gianni-Barrera et al. 2019) no increase in Piezo1 activity upon Yoda1 application was detected in cardiac MΦ post-cryoablation. This suggests either that MΦ_MD_, once recruited into cardiac tissue, undergo a phenotypic change in the direction of MΦ_TR_, or that the protocols we and others use to derive MΦ_BM_ from bone marrow (using 7-day-exposure to L929-cell conditioned medium) yields cells that differ from cardiac MΦ *in vivo*. Future experiments are needed to further characterise the molecular identity of channels underlying stretch-induced currents, and their functions in cardiac MΦ_TR_ and MΦ_MD_.

## 5. Acknowledgements

We thank all members of the Institute for Experimental Cardiovascular Medicine (IEKM) for critical discussion and input on the manuscript. We acknowledge the microscopy facility SCI-MED (Super-Resolution Confocal/Multiphoton Imaging for Multiparametric Experimental Designs) at IEKM, Freiburg, for providing access to imaging setups and analysis work stations, and its director, Josef Madl, for expert technical support. We thank the Lighthouse Facility of the University of Freiburg for providing access and support for cell sorting. All authors are members of the German Research Foundation (DFG)-funded Collaborative Research Center CRC1425 (DFG ID #422681845). The project received additional financial support *via* the Emmy-Noether program (DFG ID #412853334 to FSW) and *via* a ‘la Caixa’ Foundation PhD Fellowship to ASC (ID #100010434). AK is supported by the Berta-Ottenstein-Programme for Clinician Scientists by the Faculty of Medicine, University of Freiburg. RE is supported by a post-doctoral Fellowship from the German Cardiac Society (DGK).

## Supplementary figures

**Figure suppl 1:**
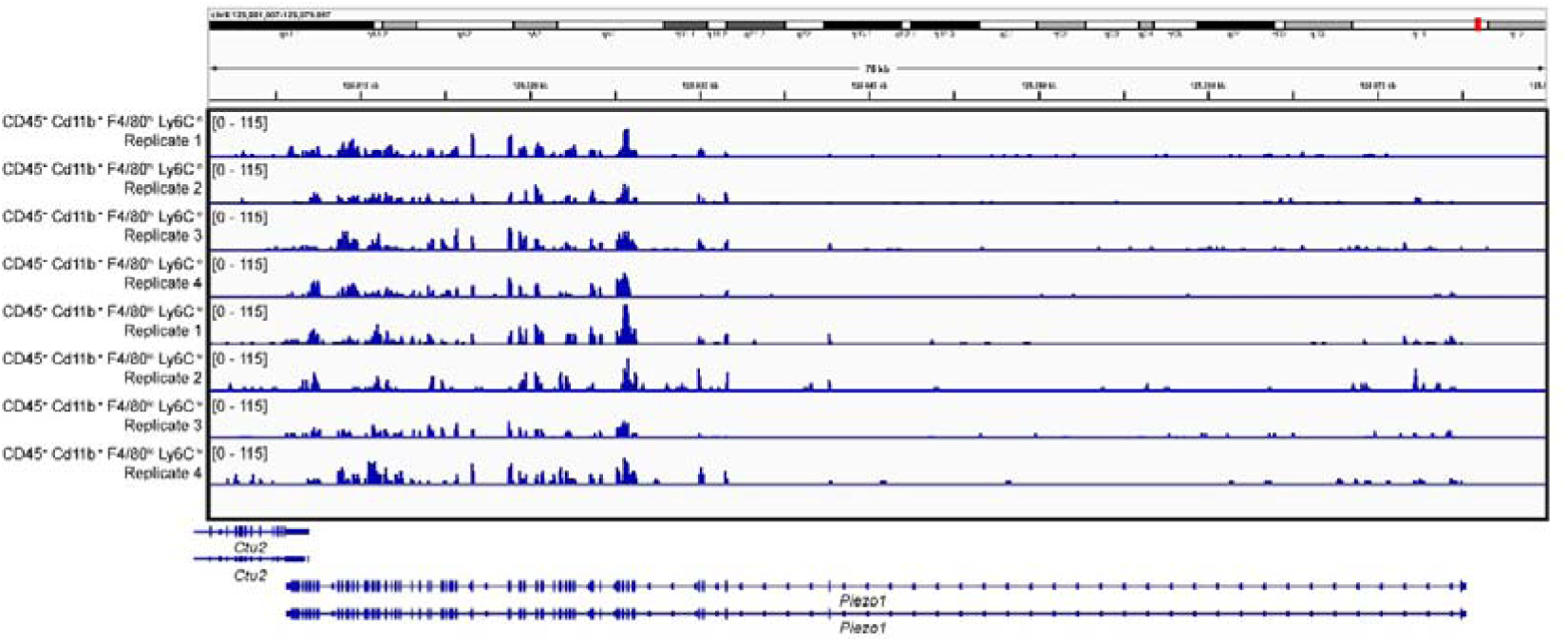
Representative traces showing the expression of *PIEZO1* in isolated cardiac MΦ_TR_ all expressing CD45.2^+^ and CD11b^+^ markers together with high F4/80 and low Ly6C (macrophages) and *vice versa (*low F4/80 and high Ly6C, monocytes*) from Lother et al.,(Lother, Bondareva et al. 2021)*.

**Figure suppl 2:**
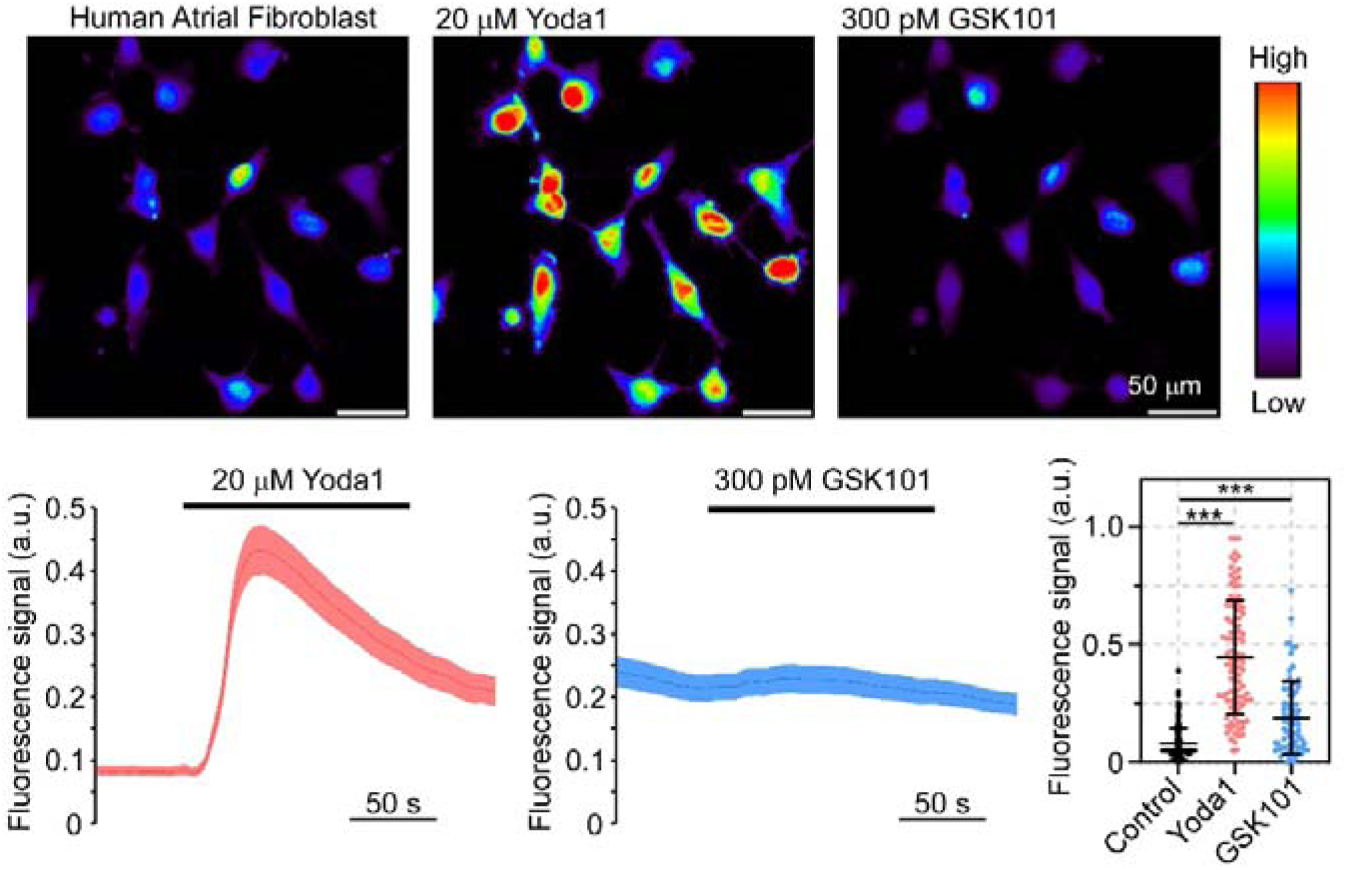
Regulation of Ca^2+^ influx in human fibroblasts upon application of Piezo1 and TRPV4 activators. Representative images showing Ca^2+^ responses to 20 μM Yoda1 and 300 μM GSK101 in human atrial fibroblasts (top). Fluorescence signal traces averaged across all cells over time and quantification of peak intensities per cell (n=151, 77 and 77 cells exposed to control, 20 μM Yoda1 and 300 μM GSK101, respectively) (bottom). Shaded regions represent SEM. Yoda1 is represented in red and GSK101 in blue. ***p < 0.001 comparing Yoda1-exposed cells with control cells and ***p < 0.001 (Student t-test).

## Bibliography

Afgan, E., D. Baker, B. Batut, M. van den Beek, D. Bouvier, M. Cech, J. Chilton, D. Clements, N. Coraor, B. A. Gruning, A. Guerler, J. Hillman-Jackson, S. Hiltemann, V. Jalili, H. Rasche, N. Soranzo, J. Goecks, J. Taylor, A. Nekrutenko and D. Blankenberg (2018). “The Galaxy platform for accessible, reproducible and collaborative biomedical analyses: 2018 update.” Nucleic Acids Res 46(W1): W537–W544.

Atcha, H., A. Jairaman, J. R. Holt, V. S. Meli, R. R. Nagalla, P. K. Veerasubramanian, K. T. Brumm, H. E. Lim, S. Othy, M. D. Cahalan, M. M. Pathak and W. F. Liu (2021). “Mechanically activated ion channel Piezo1 modulates macrophage polarization and stiffness sensing.” Nat Commun 12(1): 3256.

Bajpai, G., C. Schneider, N. Wong, A. Bredemeyer, M. Hulsmans, M. Nahrendorf, S. Epelman, D. Kreisel, Y. Liu, A. Itoh, T. S. Shankar, C. H. Selzman, S. G. Drakos and K. J. Lavine (2018). “The human heart contains distinct macrophage subsets with divergent origins and functions.” Nat Med 24(8): 1234–1245.

Baratchi, S., M. T. K. Zaldivia, M. Wallert, J. Loseff-Silver, S. Al-Aryahi, J. Zamani, P. Thurgood, A. Salim, N. M. Htun, D. Stub, P. Vahidi, S. J. Duffy, A. Walton, T. H. Nguyen, A. Jaworowski, K. Khoshmanesh and K. Peter (2020). “Transcatheter Aortic Valve Implantation Represents an Anti-Inflammatory Therapy Via Reduction of Shear Stress-Induced, Piezo-1-Mediated Monocyte Activation.” Circulation 142(11): 1092–1105.

Beaulieu-Laroche, L., M. Christin, A. Donoghue, F. Agosti, N. Yousefpour, H. Petitjean, A. Davidova, C. Stanton, U. Khan, C. Dietz, E. Faure, T. Fatima, A. MacPherson, S. Mouchbahani-Constance, D. G. Bisson, L. Haglund, J. A. Ouellet, L. S. Stone, J. Samson, M. J. Smith, K. Ask, A. Ribeiro-da-Silva, R. Blunck, K. Poole, E. Bourinet and R. Sharif-Naeini (2020). “TACAN Is an Ion Channel Involved in Sensing Mechanical Pain.” Cell 180(5): 956–967 e917.

Beech, D. J. and A. C. Kalli (2019). “Force Sensing by Piezo Channels in Cardiovascular Health and Disease.” Arterioscler Thromb Vasc Biol 39(11): 2228–2239.

Chakarov, S., H. Y. Lim, L. Tan, S. Y. Lim, P. See, J. Lum, X. M. Zhang, S. Foo, S. Nakamizo, K. Duan, W. T. Kong, R. Gentek, A. Balachander, D. Carbajo, C. Bleriot, B. Malleret, J. K. C. Tam, S. Baig, M. Shabeer, S. E. S. Toh, A. Schlitzer, A. Larbi, T. Marichal, B. Malissen, J. Chen, M. Poidinger, K. Kabashima, M. Bajenoff, L. G. Ng, V. Angeli and F. Ginhoux (2019). “Two distinct interstitial macrophage populations coexist across tissues in specific subtissular niches.” Science 363(6432).

Chen, X., S. Wanggou, A. Bodalia, M. Zhu, W. Dong, J. J. Fan, W. C. Yin, H. K. Min, M. Hu, D. Draghici, W. Dou, F. Li, F. J. Coutinho, H. Whetstone, M. M. Kushida, P. B. Dirks, Y. Song, C. C. Hui, Y. Sun, L. Y. Wang, X. Li and X. Huang (2018). “A Feedforward Mechanism Mediated by Mechanosensitive Ion Channel PIEZO1 and Tissue Mechanics Promotes Glioma Aggression.” Neuron 100(4): 799–815.e797.

Delmas, P., T. Parpaite and B. Coste (2022). “PIEZO channels and newcomers in the mammalian mechanosensitive ion channel family.” Neuron 110(17): 2713–2727.

Dick, S. A., J. A. Macklin, S. Nejat, A. Momen, X. Clemente-Casares, M. G. Althagafi, J. Chen, C. Kantores, S. Hosseinzadeh and L. Aronoff (2019). “Self-renewing resident cardiac macrophages limit adverse remodeling following myocardial infarction.” Nature immunology 20(1): 29–39.

Dutta, B., R. Goswami and S. O. Rahaman (2020). “TRPV4 Plays a Role in Matrix Stiffness-Induced Macrophage Polarization.” Front Immunol 11: 570195.

Emig, R., C. M. Zgierski-Johnston, V. Timmermann, A. J. Taberner, M. P. Nash, P. Kohl and R. Peyronnet (2021). “Passive myocardial mechanical properties: meaning, measurement, models.” Biophys Rev 13(5): 587–610.

Fernandez, M. C., R. A. Kopton, A. Simon-Chica, J. Madl, I. Hilgendorf, C. M. Zgierski-Johnston and F. Schneider-Warme (2021). “Channelrhodopsins for Cell-Type Specific Illumination of Cardiac Electrophysiology.” Methods Mol Biol 2191: 287–307.

Forget, A., R. Gianni-Barrera, A. Uccelli, M. Sarem, E. Kohler, B. Fogli, M. G. Muraro, S. Bichet, K. Aumann, A. Banfi and V. P. Shastri (2019). “Mechanically Defined Microenvironment Promotes Stabilization of Microvasculature, Which Correlates with the Enrichment of a Novel Piezo-1(+) Population of Circulating CD11b(+) /CD115(+) Monocytes.” Adv Mater 31(21): e1808050.

Gannier, F., E. White, A. Lacampagne, D. Garnier and J.-Y. L. Guennec (1994). “Streptomycin reverses a large stretch induced increase in [Ca2+]i in isolated guinea pig ventricular myocytes.” Cardiovascular Research 28(8): 1193–1198.

Heidt, T., G. Courties, P. Dutta, H. B. Sager, M. Sebas, Y. Iwamoto, Y. Sun, N. Da Silva, P. Panizzi, A. M. van der Laan, F. K. Swirski, R. Weissleder and M. Nahrendorf (2014). “Differential contribution of monocytes to heart macrophages in steady-state and after myocardial infarction.” Circ Res 115(2): 284–295.

Hulsmans, M., S. Clauss, L. Xiao, A. D. Aguirre, K. R. King, A. Hanley, W. J. Hucker, E. M. Wulfers, G. Seemann, G. Courties, Y. Iwamoto, Y. Sun, A. J. Savol, H. B. Sager, K. J. Lavine, G. A. Fishbein, D. E. Capen, N. Da Silva, L. Miquerol, H. Wakimoto, C. E. Seidman, J. G. Seidman, R. I. Sadreyev, K. Naxerova, R. N. Mitchell, D. Brown, P. Libby, R. Weissleder, F. K. Swirski, P. Kohl, C. Vinegoni, D. J. Milan, P. T. Ellinor and M. Nahrendorf (2017). “Macrophages Facilitate Electrical Conduction in the Heart.” Cell 169(3): 510–522 e520.

Jakob, D., A. Klesen, B. Allegrini, E. Darkow, D. Aria, R. Emig, A. S. Chica, E. A. Rog-Zielinska, T. Guth, F. Beyersdorf, F. A. Kari, S. Proksch, S. N. Hatem, M. Karck, S. R. Kunzel, H. Guizouarn, C. Schmidt, P. Kohl, U. Ravens and R. Peyronnet (2021). “Piezo1 and BK(Ca) channels in human atrial fibroblasts: Interplay and remodelling in atrial fibrillation.” J Mol Cell Cardiol 158: 49–62.

Lee, M., H. Du, D. A. Winer, X. Clemente-Casares and S. Tsai (2022). “Mechanosensing in macrophages and dendritic cells in steady-state and disease.” Front Cell Dev Biol 10: 1044729.

Li, M., X. Zhang, M. Wang, Y. Wang, J. Qian, X. Xing, Z. Wang, Y. You, K. Guo, J. Chen, D. Gao, Y. Zhao, L. Zhang, R. Chen, J. Cui and Z. Ren (2022). “Activation of Piezo1 contributes to matrix stiffness-induced angiogenesis in hepatocellular carcinoma.” Cancer Commun (Lond) 42(11): 1162–1184.

Liao, J., W. Lu, Y. Chen, X. Duan, C. Zhang, X. Luo, Z. Lin, J. Chen, S. Liu, H. Yan, Y. Chen, H. Feng, D. Zhou, X. Chen, Z. Zhang, Q. Yang, X. Liu, H. Tang, J. Li, A. Makino, J. X. Yuan, N. Zhong, K. Yang and J. Wang (2021). “Upregulation of Piezo1 (Piezo Type Mechanosensitive Ion Channel Component 1) Enhances the Intracellular Free Calcium in Pulmonary Arterial Smooth Muscle Cells From Idiopathic Pulmonary Arterial Hypertension Patients.” Hypertension 77(6): 1974–1989.

Liao, Y., G. K. Smyth and W. Shi (2014). “featureCounts: an efficient general purpose program for assigning sequence reads to genomic features.” Bioinformatics 30(7): 923–930.

Liu, H., J. Hu, Q. Zheng, X. Feng, F. Zhan, X. Wang, G. Xu and F. Hua (2022). “Piezo1 Channels as Force Sensors in Mechanical Force-Related Chronic Inflammation.” Front Immunol 13: 816149.

Lother, A., O. Bondareva, A. R. Saadatmand, L. Pollmeier, C. Hardtner, I. Hilgendorf, D. Weichenhan, V. Eckstein, C. Plass, C. Bode, J. Backs, L. Hein and R. Gilsbach (2021). “Diabetes changes gene expression but not DNA methylation in cardiac cells.” J Mol Cell Cardiol 151: 74–87.

McHugh, B. J., R. Buttery, Y. Lad, S. Banks, C. Haslett and T. Sethi (2010). “Integrin activation by Fam38A uses a novel mechanism of R-Ras targeting to the endoplasmic reticulum.” J Cell Sci 123(Pt 1): 51–61.

Mouton, A. J., K. Y. DeLeon-Pennell, O. J. Rivera Gonzalez, E. R. Flynn, T. C. Freeman, J. J. Saucerman, M. R. Garrett, Y. Ma, R. Harmancey and M. L. Lindsey (2018). “Mapping macrophage polarization over the myocardial infarction time continuum.” Basic research in cardiology 113: 1–18.

Nicolas-Avila, J. A., A. V. Lechuga-Vieco, L. Esteban-Martinez, M. Sanchez-Diaz, E. Diaz-Garcia, D. J. Santiago, A. Rubio-Ponce, J. L. Li, A. Balachander, J. A. Quintana, R. Martinez-de-Mena, B. Castejon-Vega, A. Pun-Garcia, P. G. Traves, E. Bonzon-Kulichenko, F. Garcia-Marques, L. Cusso, A. G. N. A. Gonzalez-Guerra, M. Roche-Molina, S. Martin-Salamanca, G. Crainiciuc, G. Guzman, J. Larrazabal, E. Herrero-Galan, J. Alegre-Cebollada, G. Lemke, C. V. Rothlin, L. J. Jimenez-Borreguero, G. Reyes, A. Castrillo, M. Desco, P. Munoz-Canoves, B. Ibanez, M. Torres, L. G. Ng, S. G. Priori, H. Bueno, J. Vazquez, M. D. Cordero, J. A. Bernal, J. A. Enriquez and A. Hidalgo (2020). “A Network of Macrophages Supports Mitochondrial Homeostasis in the Heart.” Cell 183(1): 94–109 e123.

Nikolaev, Y. A., C. D. Cox, P. Ridone, P. R. Rohde, J. F. Cordero-Morales, V. Vasquez, D. R. Laver and B. Martinac (2019). “Mammalian TRP ion channels are insensitive to membrane stretch.” J Cell Sci 132(23).

Peyronnet, R., J. M. Nerbonne and P. Kohl (2016). “Cardiac Mechano-Gated Ion Channels and Arrhythmias.” Circ Res 118(2): 311–329.

Rios, F. J., Z.-G. Zou, A. P. Harvey, K. Y. Harvey, R. Nosalski, P. Anyfanti, L. L. Camargo, S. Lacchini, A. G. Ryazanov and L. Ryazanova (2020). “Chanzyme TRPM7 protects against cardiovascular inflammation and fibrosis.” Cardiovascular research 116(3): 721–735.

Romani, P., L. Valcarcel-Jimenez, C. Frezza and S. Dupont (2021). “Crosstalk between mechanotransduction and metabolism.” Nat Rev Mol Cell Biol 22(1): 22–38.

Santoni, G., M. B. Morelli, C. Amantini, M. Santoni, M. Nabissi, O. Marinelli and A. Santoni (2018). ””Immuno-Transient Receptor Potential Ion Channels”: The Role in Monocyte- and Macrophage-Mediated Inflammatory Responses.” Front Immunol 9: 1273.

Scheraga, R. G., S. Abraham, K. A. Niese, B. D. Southern, L. M. Grove, R. D. Hite, C. McDonald, T. A. Hamilton and M. A. Olman (2016). “TRPV4 Mechanosensitive Ion Channel Regulates Lipopolysaccharide-Stimulated Macrophage Phagocytosis.” J Immunol 196(1): 428–436.

Schmidt, U., M. Weigert, C. Broaddus and G. Myers (2018). Cell detection with star-convex polygons. Medical Image Computing and Computer Assisted Intervention–MICCAI 2018: 21st International Conference, Granada, Spain, September 16-20, 2018, Proceedings, Part II 11, Springer.

Selezneva, A., A. J. Gibb and D. Willis (2022). “The contribution of ion channels to shaping macrophage behaviour.” Front Pharmacol 13: 970234.

Simon-Chica, A., M. C. Fernández, E. M. Wülfers, A. Lother, I. Hilgendorf, G. Seemann, U. Ravens, P. Kohl and F. Schneider-Warme (2022). “Novel insights into the electrophysiology of murine cardiac macrophages: relevance of voltage-gated potassium channels.” Cardiovascular Research 118(3): 798–813.

Solis, A. G., P. Bielecki, H. R. Steach, L. Sharma, C. C. D. Harman, S. Yun, M. R. de Zoete, J. N. Warnock, S. D. F. To, A. G. York, M. Mack, M. A. Schwartz, C. S. Dela Cruz, N. W. Palm, R. Jackson and R. A. Flavell (2019). “Mechanosensation of cyclical force by PIEZO1 is essential for innate immunity.” Nature 573(7772): 69–74.

Suchyna, T. M., V. S. Markin and F. Sachs (2009). “Biophysics and structure of the patch and the gigaseal.” Biophys J 97(3): 738–747.

Syeda, R., J. Xu, A. E. Dubin, B. Coste, J. Mathur, T. Huynh, J. Matzen, J. Lao, D. C. Tully, I. H. Engels, H. M. Petrassi, A. M. Schumacher, M. Montal, M. Bandell and A. Patapoutian (2015). “Chemical activation of the mechanotransduction channel Piezo1.” Elife 4.

Vicente, R., A. Escalada, M. Coma, G. Fuster, E. Sanchez-Tillo, C. Lopez-Iglesias, C. Soler, C. Solsona, A. Celada and A. Felipe (2003). “Differential voltage-dependent K+ channel responses during proliferation and activation in macrophages.” Journal of Biological Chemistry 278(47): 46307–46320.

Villalonga, N., M. David, J. Bielanska, R. Vicente, N. Comes, C. Valenzuela and A. Felipe (2010). “Immunomodulation of voltage-dependent K+ channels in macrophages: molecular and biophysical consequences.” J Gen Physiol 135(2): 135–147.

Walter, W., L. Alonso-Herranz, V. Trappetti, I. Crespo, M. Ibberson, M. Cedenilla, A. Karaszewska, V. Nunez, I. Xenarios and A. G. Arroyo (2018). “Deciphering the dynamic transcriptional and post-transcriptional networks of macrophages in the healthy heart and after myocardial injury.” Cell reports 23(2): 622–636.

Wong, N. R., J. Mohan, B. J. Kopecky, S. C. Guo, L. X. Du, J. Leid, G. S. Feng, I. Lokshina, O. Dmytrenko, H. Luehmann, G. Bajpai, L. Ewald, L. Bell, N. Patel, A. Bredemeyer, C. J. Weinheimer, J. M. Nigro, A. Kovacs, S. Morimoto, P. O. Bayguinov, M. R. Fisher, W. T. Stump, M. Greenberg, J. A. J. Fitzpatrick, S. Epelman, D. Kreisel, R. Sah, Y. J. Liu, H. Z. Hu and K. J. Lavine (2021). “Resident cardiac macrophages mediate adaptive myocardial remodeling.” Immunity 54(9): 2072-+.

Xu, J., C. Gao, Y. He, X. Fang, D. Sun, Z. Peng, H. Xiao, M. Sun, P. Zhang, T. Zhou, X. Yang, Y. Yu, R. Li, X. Zou, H. Shu, Y. Qiu, X. Zhou, S. Yuan, S. Yao and Y. Shang (2023). “NLRC3 expression in macrophage impairs glycolysis and host immune defense by modulating the NF-kappaB-NFAT5 complex during septic immunosuppression.” Mol Ther 31(1): 154–173.

Yu, Z.-Y., H. Gong, S. Kesteven, Y. Guo, J. Wu, J. V. Li, D. Cheng, Z. Zhou, S. E. Iismaa and X. Kaidonis (2022). “Piezo1 is the cardiac mechanosensor that initiates the cardiomyocyte hypertrophic response to pressure overload in adult mice.” Nature Cardiovascular Research 1(6): 577–591.

